## Supporting Information for "The Digital Brain Bank, an open access platform for post-mortem datasets"

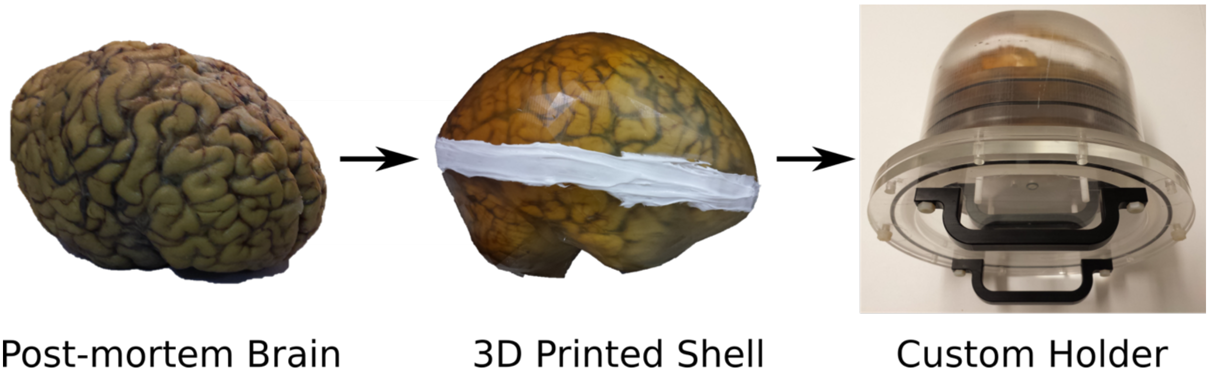

*Figure S1: Post-mortem Human Brain Holder*. The brain holder ensures consistent placement during scanning. Here, the custom holder tightly seals the brain in place, whilst the 3D printed shell (provided by Dr Alard Roebroeck, Maastricht University) prevents pressure on the brain. The holder is designed with a spherical cavity to maximise field homogeneity. For further information on our scanning procedure for human post-mortem brains, see (Wang et al., 2020)

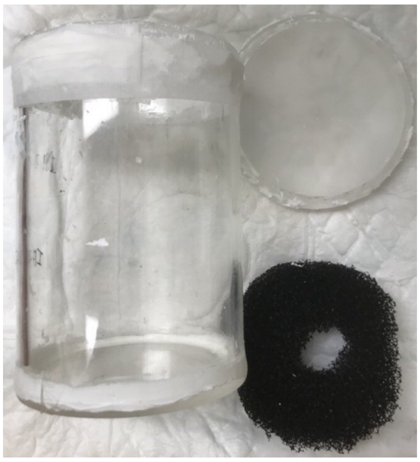

Figure S2: *Post-mortem Cylindrical Brain Holder*. The brain holder used for scanning large non-human brains which fit inside the 28 channel QED knee coil. This consisted of a cylindrical container, with plastic gauze (black) used to secure samples during the acquisition

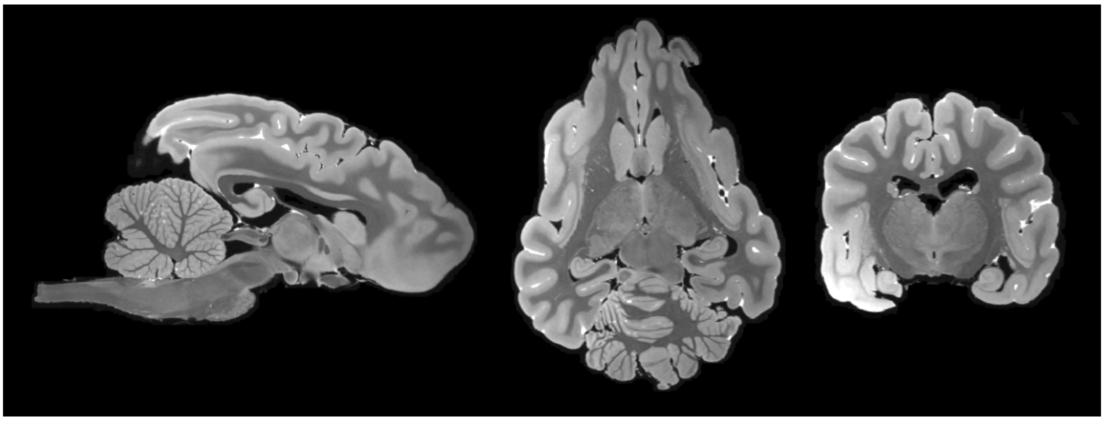

Figure S3: *Structural MRI.* Example structural MRI dataset acquired using a bSSFP sequence in the European wolf (Canis lupus) at a resolution of 220 µm (isotropic). bSSFP Structural MRI datasets display excellent grey-white matter contrast, facilitating the delineation of fine tissue structures and integration with processing pipelines for surface reconstruction. Contrast in bSSFP datasets is reversed compared to conventional T1-weighted structural MRI scans (grey matter appears bright, and white matter appears dark), which must be accounted for in any analysis pipeline.

### *Human High-Resolution Diffusion MRI-PLI dataset*

*MRI Preparation and Scanning*

Data were acquired from a post-mortem human brain (n=1) with no known neuropathology. The brain was extracted from the skull within 72 hours after death and fixed in 10% PBS buffered formalin (4% formaldehyde) for 6 weeks prior to scanning. The brain was removed from formalin and placed in plastic bags filled with Fomblin LC08 (*Solvay Solexis*), a susceptibility-matched perfluoropolyether liquid that contributes no signal to the imaging experiment.

The brain was imaged with a Siemens 7T whole body scanner (1Tx/32Rx head coil). Diffusion-weighted volumes were acquired using diffusion-weighted steady-state free precession (DW-SSFP) sequence. As highlighted in the main text, the choice of DW-SSFP was motivated by the sequences potential to simultaneously address the short $T_2$ and low diffusivity of fixed, post-mortem tissue, when limited to human scanners.

Whole-brain diffusion MRI datasets were acquired at 500 µm, 1 mm and 2 mm isotropic resolution. Details of the acquisition parameters are provided in Table S1, where we note that the 500 µm dataset took approximately 6 days of continuous scanning to acquire. DW-SSFP datasets were obtained at two flip angles to address B1-inhomogeneity at 7T, as previously described in (Tendler, Foxley, et al., 2020).

The DW-SSFP signal is dependent on tissue relaxation time-constants ($T_1$ and $T_2$) and the acquisition flip angle, which must be estimated for accurate modelling. These parameters were estimated using a turbo inversion-recovery (TIR), turbo spin-echo (TSE) and actual flip angle imaging (AFI) (Yarnykh, 2007) sequence. A structural scan was additionally acquired using a true fast imaging with steady-state precession (TRUFI) sequence (bSSFP), which produces high grey/white matter contrast in post-mortem tissue. Details of the acquisition parameters are provided in Table S2.

*MRI Processing*

A Gibbs ringing correction was applied to the DW-SSFP, TIR and TSE datasets (Kellner et al., 2016). All coregistrations within-and-between modalities were performed using a 6 degrees-of-freedom coregistration using FSL FLIRT (Jenkinson & Smith, 2001). T1 maps were estimated from the TIR volumes assuming mono-exponential signal evolution. T2 maps were estimated from the TSE volumes using an extended phase graph (EPG) framework (Weigel, 2015), as described in (Tendler, Qi, et al., 2020). B1 maps were estimated from the AFI volumes as described in the original AFI publication (Yarnykh, 2007). Structural scans were estimated from the TRUFI volumes, with banding artefacts minimized by taking the maximum intensity across volumes (Bangerter et al., 2004).

For the diffusion outputs, Tensor and Ball & 2-Stick models were fit to the DW-SSFP data as described in (Tendler, Foxley, et al., 2020). In brief, fitting was performed using the full DW-SSFP Buxton model (Buxton, 1993), estimating a shared set of diffusion orientations (e.g. tensor eigenvectors), and a unique set of diffusivity estimates (e.g. tensor eigenvalues) per DW-SSFP flip angle. The fitting process incorporated the estimated T1, T2 and B1 maps, in addition to a noise-floor correction.

The DW-SSFP sequence does not have a well-defined b-value. To address this, the diffusivity estimates at each flip angle were combined to generate diffusivity estimates at an effective b-value of 4000 s/mm$^2$. Details of this procedure, in addition to the motivation behind the choice of 4000 s/mm$^2$ are detailed in (Tendler, Foxley, et al., 2020). Note that a small modification was made to the original minimization procedure, as described in the Appendix below.

*PLI preparation, Scanning and Processing*

Tissue samples from the anterior commissure, corpus callosum, occipital lobe gurus, pons, thalamus and external capsule were extracted from the post-mortem brain. Samples were stored in a 30% sucrose solution with phosphate buffered saline (PBS) and 0.025% azide for three weeks. Tissue blocks were subsequently

| DW-SSFP (0.5 mm) | | DW-SSFP (1.0 mm) | |
|---|---|---|---|
| q-value ($cm^{-1}$) | 300 | q-value ($cm^{-1}$) | 300 |
| Diffusion Gradient Duration (ms) | 14.10 | Diffusion Gradient Duration (ms) | 14.10 |
| Diffusion Gradient Strength ($mTm^{-1}$) | 50 | Diffusion Gradient Strength ($mTm^{-1}$) | 50 |
| Flip angles (°) | 33 and 98 | Flip angles (°) | 33 and 98 |
| No. directions (per flip angle) | 90 | No. directions (per flip angle) | 60 |
| No. non-DW (per flip angle) | 6 (q=20 $cm^{-1}$) | No. non-DW (per flip angle) | 5 (q=20 $cm^{-1}$) |
| Resolution ($\mu m^3$) | 500·500·500 | Resolution ($mm^3$) | 1.0·1.0·1.0 |
| TE (ms) | 21 | TE (ms) | 21 |
| TR (ms) | 30 | TR (ms) | 30 |
| EPI factor | 1 | EPI factor | 1 |
| Bandwidth (Hz per pixel) | 198 | Bandwidth (Hz per pixel) | 130 |
| Acquisition time (per direction/non-DW) | 45m 03s | No. of averages | 1 |
| Acquisition time (total) | 6d 0h | | |
| No. of averages | 1 | | |

| DW-SSFP (2.0 mm) | |
|---|---|
| q-value ($cm^{-1}$) | 300 |
| Diffusion Gradient Duration (ms) | 14.10 |
| Diffusion Gradient Strength ($mTm^{-1}$) | 50 |
| Flip angles (°) | 33 and 98 |
| No. directions (per flip angle) | 221 |
| No. non-DW (per flip angle) | 6 (q=20 $cm^{-1}$) |
| Resolution ($mm^3$) | 2.0·2.0·2.0 |
| TE (ms) | 21 |
| TR (ms) | 30 |
| EPI factor | 1 |
| Bandwidth (Hz per pixel) | 130 |
| No. of averages | 1 |

Table S1: DW-SSFP Acquisition parameters at 0.5, 1.0 and 2.0 mm.

embedded in optimal cutting temperature (OCT) compound (Sakura, Finetek Inc, USA) and frozen to -80 °C. 60 µm sections were cut from the tissue blocks with a cryostat microtome (Leica, Germany). No tissue staining was performed, as birefringence is naturally expressed by the myelin sheath.

PLI was performed using a Leica DM4000B microscope, equipped with a polarizing filter, a quarter wave plate (QWP) and a rotatable analyser with orientation $\rho$. Samples were illuminated with a white LED (pE-100$^{wht}$ Cooled). The fast axis of the QWP was oriented 45° with respect to the transmission axis of the polarising filter to create circular polarization. The rotating analyser captured the phase shift induced by the myelin sheath. A total of 18 images were acquired for each field of view at equidistant analyser orientation angles, $\rho = \{0°, 10°, \ldots, 170^{\circ}\}$. Images were magnified x1.25 (0.04 NA, Leica) and captured with a Leica DFC420 CCD camera (4 µm/pixel). The green colour image channel was used for further analysis.

The entire sample was imaged via raster scanning, with each row composed of multiple contiguous field-of-views (FOV). These FOVs were automatically stitched together using in-house software (MATLAB 2015b, MathWorks, Natick, MA, USA). For each specimen, a series of background images were acquired to correct for illumination inhomogeneities (Dammers et al., 2010). Microscopic fibre orientations were derived using Jones calculus (Jones, 1941), as described in the Appendix below.

| Turbo inversion-recovery (TIR) | | True-Fast Imaging with SSFP (TRUFI) | |
|---|---|---|---|
| Resolution (mm$^3$) | 0.75·0.75·1.60 | Resolution (μm$^3$) | 312.5·312.5·500 |
| Number of inversions | 6 | TE (ms) | 5.95 |
| TE (ms) | 12 | TR (ms) | 11.9 |
| TR (ms) | 1000 | Flip angle (°) | 35 |
| TIs (ms) | 31, 62, 125, 250, 500 & 850 | Bandwidth (Hz per pixel) | 130 |
| Flip angle (°) | 180 | Phase increments (°) | 0, 180 |
| Bandwidth (Hz per pixel) | 199 | Number of averages (per set of increments) | 16 |
| Number of averages | 1 | | |
| Turbo spin-echo (TSE) – $T_2$ | | Actual flip-angle imaging (AFI) – $B_1$ | |
| Resolution (mm$^3$) | 0.75·0.75·1.60 | Resolution (mm$^3$) | 1.50·1.50·1.50 |
| Number of echoes | 6 | TE (ms) | 1.5 |
| TEs (ms) | 14, 28, 42, 56, 70 & 84 | $TR_1/TR_2$ (ms) | 6/30 |
| TR (ms) | 1000 | Flip angle (°) | 60 |
| Flip angle (°) | 180 | Bandwidth (Hz per pixel) | 630 |
| Bandwidth (Hz per pixel) | 130 | Number of averages | 1 |
| Number of averages | 1 | | |

Table S2: Acquisition parameters for the TIR, TSE, structural (TRUFI) and B1-mapping (AFI) sequences.

### *Human ALS MRI-Histology Callosum analysis*

A comparison of diffusivity properties between the ALS and control cohort (12 ALS and 3 control brains) was performed in the corpus callosum, as displayed in Figure 3c (Main Text). To achieve this, a standard-space mask of the corpus callosum was first generated using the Jülich atlas (Eickhoff et al., 2005). The callosum mask was subsequently split into five distinct regions of interest (ROI) associated with specific fibre projections as proposed by Hofer and Frahm (Hofer & Frahm, 2006), and transformed into the space of each post-mortem brain. Briefly, a standard space fractional anisotropy (FA) template (FMRIB58_FA, available as part of FSL) was modified to display similar contrast to the post-mortem FA maps. Coregistration matrices were subsequently estimated between the FA map of each post-mortem brain and the modified standard space FA template using a non-linear coregistration (ANTS) (Avants et al., 2011). The callosum masks were subsequently coregistered into the space of each post-mortem brain using the estimated coregistration matrices, and multiplied by a white matter mask (generated using FAST (Zhang et al., 2001)) to remove any remaining grey matter regions.

Diffusion estimates were obtained by taking the mean over each ROI. Values were normalised to the splenium (Par/Temp/Occ) estimate, which has been proposed as a control region with little pathological burden in ALS (Cardenas et al., 2017). Differences in the normalised FA, MD, axial and radial diffusivity between the ALS and control cohort were assessed with a two-tailed, family-wise error rate (FWER) corrected t-test using PALM (Winkler et al., 2014). Full results are provided in Table S3. Although our statistical analysis does account for sample size, it does not consider other confounds that may contribute to differences between the two groups (e.g. age, sex). Source data for the corpus callosum analysis are provided in a Supplementary File.

| | Body (Sensory) | Body (Motor) | Body (Pre/Supp Motor) | Genu (Pre-Frontal) |
|---|---|---|---|---|
| Fractional anisotropy (FA) | $p = 0.34$<br>$p_{FWER} = 0.70$ | $p = 0.042$*<br>$p_{FWER} = 0.11$ | $p = 0.0044$*<br>$p_{FWER} = 0.013$** | $p = 0.013$*<br>$p_{FWER} = 0.037$** |
| Mean Diffusivity (MD) | $p = 0.99$<br>$p_{FWER} = 1.00$ | $p = 0.18$<br>$p_{FWER} = 0.48$ | $p = 0.015$*<br>$p_{FWER} = 0.053$ | $p = 0.037$*<br>$p_{FWER} = 0.12$ |
| Axial Diffusivity (AD) | $p = 0.53$<br>$p_{FWER} = 0.92$ | $p = 0.58$<br>$p_{FWER} = 0.95$ | $p = 0.084$<br>$p_{FWER} = 0.27$ | $p = 0.11$<br>$p_{FWER} = 0.32$ |
| Radial Diffusivity (RD) | $p = 0.66$<br>$p_{FWER} = 0.97$ | $p = 0.073$<br>$p_{FWER} = 0.23$ | $p = 0.0022$*<br>$p_{FWER} = 0.015$** | $p = 0.022$*<br>$p_{FWER} = 0.062$ |

Table S3: p-values associated with differences between the ALS and control cohort for the diffusivity estimates. Here 'p' defines the p-value, and '$P_{FWER}$' defines the FWER-corrected p-value (∗= $p < 0.05$; ∗∗ = $p_{FWER} < 0.05$). The largest differences between the ALS and control cohort were found in the Body (Pre/Supp Motor) category, followed by the Genu (Pre-Frontal) and Body (Motor) category. No differences were found in the Body (Sensory) category.

### *Digital Brain Zoo Datasets*

Here we provide the acquisition and processing protocol for four previously unreleased datasets to the Digital Brain Zoo.

*European wolf and Hamadryas baboon*

Formalin fixed European wolf and Hamadryas baboon brains were provided by Copenhagen zoo. Prior to scanning, the brains were rehydrated using a phosphate-buffered saline solution. The size of the European wolf and Hamadryas baboon brain necessitated scanning on the Siemens 7T whole body scanner (1Tx/28Rx knee coil, QED), using the brain holder displayed in Supporting Information Fig. S2. The brain holder was filled with fluorinert (FC-3283, *3M*) during the scanning procedure, a susceptibility matched fluid that gives off no signal. Diffusion-weighted volumes were acquired using diffusion-weighted steady-state free precession (DW-SSFP) sequence. As highlighted in the main text, the choice of DW-SSFP was motivated by the sequences potential to simultaneously address the short $T_2$ and low diffusivity of fixed, post-mortem tissue, when limited to human scanners. A structural scan was additionally acquired using a true fast imaging with steady-state precession (TRUFI) sequence (bSSFP), which produces high grey/white matter contrast in post-mortem tissue. Acquisition parameters are provided in Supporting Information Table S3.

Structural scans were formed by averaging over all 16 TRUFI datasets (root-mean sum of squares). Diffusion datasets were processed using a similar approach to the great ape datasets in (Bryant et al., 2021) and (Roumazeilles et al., 2020). In brief, a Gibbs ringing correction (Kellner et al., 2016) was applied to the diffusion and non-diffusion weighted datasets, with all coregistrations performed using FSL FLIRT (Jenkinson & Smith, 2001). Fitting was performed using the full DW-SSFP Buxton model (Buxton, 1993) adapted to incorporate diffusion tensor and ball & 2 stick estimates. The fitting process incorporated estimated T1, T2 and B1 maps derived from a turbo inversion recovery (TIR), turbo spin-echo (TSE) and actual flip angle imaging (AFI) (Yarnykh, 2007) sequence acquired in the same session.

| DW-SSFP | | True-Fast Imaging with SSFP (TRUFI) | |
|---|---|---|---|
| q-value ($cm^{-1}$) | 300 | Resolution ($\mu m^3$) | 217·217·220 |
| Diffusion Gradient Duration (ms) | 13.56 | TE (ms) | 7.33 |
| Diffusion Gradient Strength ($mTm^{-1}$) | 52 | TR (ms) | 14.65 |
| Flip angle (°) | 39 | Flip angle (°) | 30 |
| No. directions | 160 | Bandwidth (Hz per pixel) | 100 |
| No. non-DW | 13 / 11 (q=20 $cm^{-1}$) | Phase increments (°) | 0, 45, 90, 135, 180, 225, 270, 315 |
| Resolution ($\mu m^3$) | 600·600·600 | Number of averages (per set of increments) | 2 |
| TE (ms) | 21 | | |
| TR (ms) | 29 | | |
| EPI factor | 1 | | |
| Bandwidth (Hz per pixel) | 100 | | |
| Acquisition time (per direction/non-DW) | 16m 25s | | |
| Acquisition time (total) | 1d 20h | | |
| No. of averages | 1 | | |

Table S3: Acquisition parameters for the DW-SSFP and structural (TRUFI) sequences. The only difference between the European wolf and Hamadryas baboon acquisition was the number of non-diffusion weighted directions acquired (13 for wolf, 11 for baboon).

*Cotton-Top and Golden Lion tamarins*

Formalin fixed Cotton-Top and Golden Lion tamarin brains were provided by Copenhagen zoo. Prior to scanning, the brains were rehydrated using a phosphate-buffered saline solution. Scanning was performed using a 7T magnet with Agilent Direct-Drive console and 72mm ID quadrature birdcage RF coil (Rapid Biomedical GmbH). The brain holder was filled with fluorinert during the scanning procedure, a susceptibility matched fluid that gives off no signal. Diffusion-weighted volumes were acquired using diffusion-weighted spin-echo protocol with single line readout (DW-SEMS) sequence. Acquisition parameters are provided in Supporting Information Table S4.

The diffusion datasets were processed using a similar approach to the prosimian and monkey data in (Bryant et al., 2021). Datasets were preprocessed using FSL tools implemented in the Phoenix module of the MR Comparative Anatomy Toolbox (Mr Cat, www.neuroecologylab.org). Diffusion tensor and ball & 2/3 stick estimates were derived using FSL's dtifit and bedpostX (Behrens et al., 2007).

| DW-SEMS | |
|---|---|
| b-value (s/mm$^2$) | 4000 |
| $\delta$ (ms) | 7 |
| $\Delta$ (ms) | 13 |
| Diffusion Gradient Strength (mTm$^{-1}$) | 320 |
| No. directions | 128 |
| No. non-DW | 16 |
| Resolution (µm$^3$) | 300·300·300 |
| TE (ms) | 25 |
| TR (s) | 10 |
| EPI factor | 1 |
| Bandwidth (kHz) | 100 |
| Acquisition time (per direction/non-DW) | 21m 20s |
| Acquisition time (total) | 2d 4h |
| No. of averages | 1 |

Table S4: Acquisition parameters for the DW-SEMS sequence.

### *Appendix*

*Modification to minimisation procedure*

(Tendler, Foxley, et al., 2020) described an approach to estimate DW-SSFP diffusivity estimates at a single effective b-value, achieved by incorporating a non-Gaussian diffusion model into the DW-SSFP signal equations. In (Tendler, Foxley, et al., 2020), non-Gaussianity was modelled using a Gamma distribution of diffusivities, estimating a mean ($D_m$) and standard deviation ($D_s$) of the Gamma distribution per voxel. Here, the Gamma fitting procedure (Eq. (4) in (Tendler, Foxley, et al., 2020)) was replaced with:

$$\min_{D_{m_i}, D_{s_i}} \left\| \frac{\left(L_{i_{\text{sim}:\alpha_{\text{low}}}}\left(D_{m_i}, D_{s_i}\right) - L_{i_{\text{exp}:\alpha_{\text{low}}}}\right)}{SD\left(L_{i_{\text{exp}:\alpha_{\text{low}}}}\right)} \right\|_2^2 + \left\| \frac{\left(L_{i_{\text{sim}:\alpha_{\text{high}}}}\left(D_{m_i}, D_{s_i}\right) - L_{i_{\text{exp}:\alpha_{\text{high}}}}\right)}{SD\left(L_{i_{\text{exp}:\alpha_{\text{high}}}}\right)} \right\|_2^2, \quad (1)$$

where $\alpha_{\text{low}}$ and $\alpha_{\text{high}}$ are the voxelwise DW-SSFP flip angles, $L_{i_{\text{exp}:\alpha_{\text{low}}/\alpha_{\text{high}}}}$ are the voxelwise experimental diffusivity estimates (Diffusion Tensor eigenvalues or Ball and 2-Stick diffusivity estimates) at each flip angle, $L_{i_{\text{sim}:\alpha_{\text{low}}/\alpha_{\text{high}}}}$ are the simulated diffusivity estimates for a given $D_{m_i}$ and $D_{s_i}$, and $SD\left(L_{i_{\text{exp}:\alpha_{\text{low}}/\alpha_{\text{high}}}}\right)$ are the estimated experimental standard deviation of the diffusivity estimates. This approach was found to reduce spurious diffusivity estimates in regions of low SNR. For further details of the modelling approach, see (Tendler, Foxley, et al., 2020).

*PLI Fibre Orientations*

The light intensity ($I$) for a birefringent specimen inside a PLI-setup is described using Jones calculus (Jones 1941), defining:

$$I(\rho) = \frac{I_0}{2}[1 + \sin(2\rho - 2\varphi) \cdot \sin\delta], \quad (1)$$

where $I_0$ is the average light intensity, $\rho$ the polarizer orientation, $\varphi$ is the in-plane orientation of the myelin sheath and $\delta$ is the phase shift, defined as:

$$\delta \approx 2\pi \frac{d \cdot \Delta n}{\lambda} \cdot \cos^2\alpha, \quad (2)$$

where $d$ is the sample thickness, $\Delta n$ is the sample birefringence, $\lambda$ is the light wavelength and $\alpha$ is the inclination angle of the myelin sheath (α). Microscopic fibre orientations were derived using as above, fitting to each pixel in the raw PLI images as previously reported in (Axer et al., 2011).
