## Supplementary material for "The Digital Brain Bank, an open access platform for post-mortem datasets": README.txt

Data used to perform the ALS vs Control corpus callosum analysis in Figure 3c

Files are as follows:

Axial.csv	- Axial diffusivity values for the ALS and Control brains over the Body (Sensory), Body (Motor), Body (Pre/Supp-Motor) and Genu (Pre-Frontal) regions, normalised by the Splenium (Par/Temp/Occ) value.
FA.csv 		- Fractional Anisotropy values for the ALS and Control brains over the Body (Sensory), Body (Motor), Body (Pre/Supp-Motor) and Genu (Pre-Frontal) regions, normalised by the Splenium (Par/Temp/Occ) value.
MD.csv		- Mean Diffusivity values for the ALS and Control brains over the Body (Sensory), Body (Motor), Body (Pre/Supp-Motor) and Genu (Pre-Frontal) regions, normalised by the Splenium (Par/Temp/Occ) value.
Radial.csv	- Radial Diffusivity values for the ALS and Control brains over the Body (Sensory), Body (Motor), Body (Pre/Supp-Motor) and Genu (Pre-Frontal) regions, normalised by the Splenium (Par/Temp/Occ) value.

Contrast.csv	- Contrast matrix
Design.csv 	- Design matrix (12 ALS estimates and 3 Control estimates)

Config.txt	- Palm configuration file
